# Microbubble-Enhanced Focused Ultrasound Improves Targeted Adeno-Associated Virus Delivery in Brain Tumors Quantified by PET Imaging

**DOI:** 10.64898/2026.02.06.704523

**Authors:** Yutong Guo, Josquin Foiret, Jai Woong Seo, Nisi Zhang, James Wang, Marina N. Raie, Basit L. Jan, Spencer K. Tumbale, Katherine W. Ferrara

**Affiliations:** Molecular Imaging Program at Stanford (MIPS), Department of Radiology, School of Medicine, Stanford University, Stanford, CA, USA

**Keywords:** Image-guided delivery, Adeno-associated virus, Microbubble-enhanced focused ultrasound, Positron emission tomography, Blood-tumor barrier, Glioblastoma

## Abstract

Gene therapy using adeno-associated virus (AAV) vectors shows promise for cancer treatment through molecular intervention, yet achieving sufficient and targeted delivery to brain tumors via systemic administration remains limited by the biological barriers. Here, we investigate whether microbubble-enhanced focused ultrasound (MB-FUS) improves targeted delivery of systemically administered AAV9 to orthotopic gliomas, using quantitative PET imaging of ^64^Cu-radiolabeled AAV9 vectors and fluorescent reporter expression to assess biodistribution and functional efficacy. At 21 hours after injection, ^64^Cu-AAV9 accumulation was 3.2-fold higher in FUS-treated tumors compared to non-FUS-treated tumors (n=3, p=0.004). Quantitative PCR analysis of tumor tissue at the same timepoint confirmed a 6.4-fold increase in genome copies in FUS-treated tumors (p=0.0003). The enhanced vector delivery translated to a 5.3-fold increase in optical reporter protein expression in FUS-treated compared to control tumors (p=0.0002) at 17 days post-treatment. These results establish that MB-FUS enables spatially-targeted AAV delivery with quantifiable enhancement in both acute vector biodistribution and downstream transgene expression. The integration of radiolabeled AAV with PET imaging provides a non-invasive methodology for real-time assessment of vector delivery and optimization of treatment protocol for brain cancer gene therapy.

**Highlights:**

- MB-FUS enables targeted systemic AAV delivery to brain tumors.
- MB-FUS enhanced vector delivery translates to increased transgene expression in gliomas.
- PET imaging of radiolabeled AAV allows non-invasive tracking of gene therapy vectors.
- Real-time imaging validates spatially-controlled gene delivery for brain cancer.

## Introduction

Despite extensive research and advances in therapy, from multimodal regimens combining surgery, radiotherapy, and chemotherapy to emerging immunotherapies and targeted molecular agents, malignant brain tumors remain among the most devastating cancer diagnoses [1–5]. Glioblastoma (GBM), the most aggressive subtype, has a median survival of only 15 months, with recurrent disease reducing survival to less than one year [6,7]. These treatment strategies, which have shown promise in other cancer types, face formidable obstacles in brain tumors: infiltrative growth that extends beyond resectable margins, intrinsic chemo-radio resistance, immunosuppressive microenvironments, and restrictive biological barriers that limit drug delivery [5,8,9]. Overcoming these barriers requires both novel therapeutic agents and effective delivery strategies.

Gene therapy using adeno-associated viruses (AAVs) represents a promising approach to target multiple mechanisms of treatment resistance [10,11]. By transducing tumor cells and cancer-associated stromal cells, AAVs enable sustained expression of diverse therapeutic payloads, including tumor suppressors [12], pro-apoptotic factors [13,14], anti-angiogenic agents [15–17], and immunostimulatory cytokines [18–22], offering versatility not achievable with conventional therapies. However, realizing the therapeutic potential of AAV gene therapy for brain tumors faces a critical obstacle: achieving spatially targeted vector delivery across the blood-brain and blood-tumor barriers (BBB/BTB) and adequate penetration and distribution within the tumor microenvironment [23].

Microbubble-enhanced focused ultrasound (MB-FUS) is emerging as a non-invasive tool for overcoming delivery barriers in brain tumors [24,25]. By precisely targeting the tumor regions, MB-FUS transiently increases BBB/BTB permeability and enhances interstitial transport, creating a therapeutic window for enhanced drug delivery [24,26,27]. This approach has been leveraged in completed and ongoing clinical trials to enhance delivery of a variety of therapeutics, including chemotherapeutics and antibodies, to brain tumors, with demonstrated safety and preliminary evidence of efficacy [28]. Despite these promising capabilities and growing clinical application, the potential of MB-FUS to enhance AAV vector delivery and biodistribution in brain tumors has not been investigated. While MB-FUS has been shown to enhance AAV delivery in healthy brain tissue [29–41], the heterogeneous tumor microenvironment may significantly alter vector distribution and transduction efficiency [9]. Furthermore, current assessment methods such as gadolinium-based contrast magnetic resonance imaging (MRI), while useful for confirming BBB opening, do not provide quantitative information on AAV biodistribution or transduction [42]. Quantitative and longitudinal imaging is therefore needed to directly measure AAV biodistribution kinetics with MB-FUS and assess improvements in tumor delivery.

In this work, we investigated whether MB-FUS can enhance spatially targeted AAV delivery to orthotopic GL26 glioma model using quantitative imaging to assess vector biodistribution and transgene expression in vivo. We radiolabeled AAV9 vectors with ^64^Cu and encoded a fluorescent reporter gene to enable dual-modality assessment: (1) non-invasive, longitudinal positron emission tomography (PET) imaging to track biodistribution at multiple time points following systemic administration and MB-FUS treatment, and (2) optical imaging to assess biodistribution of transgene expression with fluorescence microscopy for cellular-level analysis. Our approach provides direct, quantitative assessment of both MB-FUS-mediated AAV delivery and functional transduction in brain tumors, establishing a framework for optimizing the treatment protocol to maximize therapeutic effect.

## Methods

### In vivo experiments

All animal experiments were conducted with a protocol approved by the Institutional Animal Care & Use Committee (IACUC) at Stanford University. 2-3 months old B6 albino female mice were used in this study (Jackson Laboratory).

### Gl26 glioma cells and tumor inoculation

The murine glioma cell line GL26, genetically modified to express firefly luciferase and eGFP, was provided by Sanjiv Sam Gambhir’s laboratory at Stanford University. GL26 glioma cells were cultured in DMEM media with 10% FBS, 1% Penicillin-Streptomycin (Gibco). 10^5^ GL26 cells were stereotactically implanted into the brain at 1 mm anterior and 1 mm to the right of the Bregma of 2 month-old mice. After cell implantation, tumor growth was monitored using bioluminescence imaging and T2 weighted Magnetic Resonance Imaging (MRI) (Pharmascan 7T, Bruker). To minimize differences in baseline BBB permeability across experimental groups due to tumor size variability, mice were stratified by tumor volume and evenly distributed between non-focused ultrasound (FUS) and FUS-treated groups before treatment.

### Microbubble preparation and characterization

The microbubbles (MBs) used in this study were lipid-shelled MBs produced in-house (distearoylphosphatidylcholine (DSPC):1,2distearoyl-sn-glycero-3-phosphoethanolamine-N-[ma leimide (polyethylene glycol)-2000 (DSPE-PEG2000); 90:10 molecular ratio). To ensure robust and consistent BBB/BTB opening, we utilized monodisperse MBs with a 5-μm diameter. These MBs were isolated through size-selective centrifugation of freshly activated polydisperse solution [43]. To make sure the MB sizes and number of MBs selected were consistent using this method, the selected MBs were counted and characterized by an AccuSizer 770A Optical Particle Sizer (Santa Barbara, CA, USA) in triplicate before every MB-FUS treatment.

### Focused Ultrasound System and Treatment

We employed an ultrasound-guided focused ultrasound (USgFUS) system (1.5 MHz) featuring a 128-element 2D therapeutic array integrated with an L12-5 imaging transducer (Philips/ATL), as previously described [44]. A Verasonics Vantage 256 system was used to achieve controlled image guidance, therapy delivery, and acoustic monitoring. The therapeutic transducer was calibrated transcranially using a needle hydrophone (HNP0400, Onda) with age-matched mouse skulls positioned to simulate tumor targeting geometry (focal depth ∼4 mm). To ensure complete tumor coverage, the focal zone was raster-scanned across a 5 × 5 grid with 0.5 mm non-overlapping spacing.

Tumor-bearing mice were anesthetized and positioned supine with stereotaxic head fixation. Tumors were localized using real-time ultrasound imaging confirmed with pre-acquired MRI. AAV vectors or fluorescently labeled dextran were administered intravenously 1 minute before sonication. Treatment parameters were: 1 ms pulses, 5 Hz repetition rate, 320 kPa peak negative focal pressure, 2 minutes total duration, with concurrent intravenous microbubble administration (2.5 × 10^8^ MBs/kg). Control animals (non-FUS group) received the same microbubble dose and were anesthetized for equivalent duration to isolate the effect of sonication. Real-time passive acoustic mapping (PAM) with the angular spectrum method [45] via the imaging transducer was employed to localize acoustic emissions and characterize microbubble cavitation dynamics throughout sonication. For the analysis of the cavitation signals, the power levels for the different frequency bands were calculated as follow: the harmonics band was the 3^rd^ to 7^th^ harmonics(i.e. 4.5, 6.0,…,10.5 MHz) with a 0.1 MHz window, the ultra-harmonics band was 5.25, 6.75, 8.25, 9.75 MHz with a 0.1 MHz window, and broadband was all the other frequencies between 7.1 and 10.4 MHz.

### Magnetic resonance imaging

MRI was performed using a Bruker 11.7 Tesla small animal scanner (Bruker BioSpin MRI, Ettlingen, Germany) equipped with a cross coil configuration with a mouse body resonator for transmission and a mouse surface coil for reception. Images were acquired using ParaVision 360 (Bruker BioSpin MRI). Tumor size was assessed using a T2-weighted MRI sequence (2D FLASH sequence, flip angle (FA) = 15°, repetition time (TR) 250 ms, echo time (TE) 15 ms, 1 mm slice thickness, 1 mm interslice distance, 13 images, field of view (FOV) = 2 × 2 cm^2^, matrix = 192 × 192, number of acquisitions (NA) = 4). To confirm the BBB opening, immediately after the sonication, the animals were injected with the gadolinium contrast agent, Gd-HPDO3A (Prohance®, 0.5 μmol/g mouse body weight). Permeability of the BBB was determined with a T1 weighted (T1w) sequence (2D RARE sequence, RARE factor = 2, repetition time (TR) 250 ms, echo time (TE) 6.7 ms, 1 mm slice thickness, 1 mm interslice distance, 13 images, field of view (FOV) = 2 × 2 cm, matrix = 384 × 384, number of acquisitions (NA) = 6).

### Fluorescent optical imaging

Tumor growth, dextran delivery, and transgene expression were monitored using the LAGO imaging system (Spectral Instruments Imaging, Tucson, AZ). For bioluminescence imaging of Fluc-expressing GL26 tumors, mice received intraperitoneal D-luciferin (150 mg/kg) and were imaged 10 minutes later with 2-second exposure. For fluorescence imaging, the appropriate filter sets were used: excitation: 535 nm, emission: 589 nm. Both in vivo and ex vivo fluorescence images were acquired with 2-second exposure. Regions of interest (ROIs) were drawn over tumors or organs, and signal was quantified as total flux (photons/second) for bioluminescence or radiance/total emission for fluorescence using Aura software (Spectral Instruments).

### AAV production

AAV9 vectors were produced by the University of North Carolina Vector Core. Briefly, AAVs were harvested 5 days after triple transfection in HEK293 cells by PEG precipitation of 3- and 5-days media and osmotic lysis of cell pellets. Crude AAVs were then purified by extraction from iodixanol density gradients and buffer exchanged into Dulbecco’s phosphate-buffered saline (DPBS). Viral titers were determined by quantitative polymerase chain reaction (qPCR) on a woodchuck hepatitis virus post-transcriptional regulatory element (WPRE) present in all packaged AAV genomes.

### Radiolabeling of AAV

All radiolabeling experiments were conducted under the Controlled Radiation Authorization (CRA) approved by Stanford University (Palo Alto, CA). Radiolabeling of AAVs was performed using our previously reported method.^43,46^ [^64^Cu]CuCl_2_ was produced at the MIR Cyclotron Facility at Washington University School of Medicine. The buffer (350 mM NaCl and 5% D-sorbitol in PBS) of AAV9 was exchanged to 1XPBS in mini dialysis device (Thermo Scientific, 20k MWCO) overnight. The AAV9 (1.4×10^13^ vg) in 1xPBS (0.9 mL) was mixed with aqueous Na_2_CO_3_ (0.1 M, 60 μL, pH = 9.2), 2 mM tetrazine-PEG_5_-NHS (Tz-PEG_5_-NHS, 60 nmol, 30 μL in DMSO) and incubated at 25ºC for 30 minutes. After addition of PBS (0.1 mL), the mixture was transferred to a mini-dialysis device (20 kDa MWCO), dialyzed in PBS (15 mL) for 4 hours and subsequently transferred for overnight dialysis (500 mL).

For the radiolabeling of dialyzed Tz-AAV9, 10 μM (NOTA)_8_-TCO (40-100 pmol, 4-10 μL) in ammonium citrate buffer (20 μL, pH 6.5) was incubated with [^64^Cu]CuCl_2_ (4 mCi), 5 μL) for 30 minutes at room temperature. The incorporation of ^64^Cu to (NOTA)_8_-TCO was completed in 30 minutes as monitored by instant thin-layer chromatography (iTLC). The (^64^Cu-NOTA)_8_-TCO solution was mixed with dialyzed Tz-AAV9s and incubated for 30 min. The radiolabeled AAV9 mixture was further purified by three cycles of dilution and concentration with PBS (3x15 mL) containing 0.001% Pluronic F-68 (Gibco) in a 100 kDa MWCO centrifugal filter unit (Thermo Fisher Scientific). ^64^Cu-AAV9 (54 μCi) was recovered in 200 μL volume of PBS.

### PET imaging

Longitudinal PET/CT imaging was performed at 0.5, 6, and 21 hours post-treatment using an Inveon microPET/CT scanner (Siemens Medical Solutions). Mice were anesthetized with isoflurane (1-2% in oxygen) and positioned prone with the head centered in the field of view. Static PET acquisitions were performed for 30 minutes per timepoint, followed by CT imaging for anatomical co-registration.

PET data were reconstructed using 3D ordered-subset expectation maximization with a maximum a posteriori (3D-OSEM/MAP) algorithm and corrected for decay, dead time, and scatter. PET and CT images were co-registered using Inveon Research Workplace (IRW) software (Siemens). Regions of interest (ROIs) were manually drawn over tumors and contralateral non-treated brain region. Activity concentration was quantified as percent injected dose per cubic centimeter (%ID/cc). For longitudinal comparison, values were decay-corrected to the time of injection.

### DNA extraction

To assess AAV9 vector biodistribution, mice were euthanized 21 hours post-treatment, and brains were extracted and bisected on ice. Tumor tissue from the FUS-treated hemisphere and non-tumor tissue of similar volume from the contralateral hemisphere were dissected. The brain sample was immediately stabilized in Allprotect Tissue Reagent (Qiagen) until processing. DNA was isolated from tissue homogenates using the DNeasy Blood & Tissue Kit (Qiagen) according to the manufacturer’s protocol. DNA concentration and purity were determined by NanoDrop spectrophotometry (Thermo Fisher Scientific), and samples were diluted to a uniform concentration for qPCR analysis.

### RNA extraction and cDNA synthesis

To assess transgene expression, brain tissue was collected and immediately stabilized in RNA-protect tissue reagent (Qiagen) on ice. Total RNA was extracted from brain hemispheres using the RNeasy Midi Kit (Qiagen) following the manufacturer’s instructions. During this process, genomic DNA was removed by DNase. RNA integrity and concentration were assessed by NanoDrop spectrophotometry, and samples with an A260/A280 ratio between 1.8 and 2.0 were used for downstream analysis. Complementary DNA (cDNA) was synthesized from 500 ng - 1 µg of total RNA using SuperScript IV VILO Master mix (ThermoFisher Scientific) according to the manufacturer’s protocol.

### Quantitative PCR (qPCR)

qPCR was performed using TaqMan-based assays with primers and probe designed to detect specific genome copies (WPRE, tdTomato) and normalized to the housekeeping gene, β-actin. Relative expression was calculated using the ΔCt method and expressed as fold-change relative to control groups. All reactions were performed in triplicate on a CFX96 (Bio-Rad) system using the following cycling conditions: 95°C for 10 min, followed by 40 cycles of 95°C for 15 s and 60°C for 1 min.

### Immunofluorescence staining and microscopy

For protein expression analysis, after the animals were euthanized at the time point based on their treatment protocols, the brains were harvested and were fixed with 4% paraformaldehyde overnight at 4°C. The next day, 20 µm sections were cut using a microtome (Leica 3050 S Cryostat). The sections were imaged with a 20x objective using a fluorescence confocal microscope (Leica DMi8 Inverted Microscope). The quantification of the fluorescence images was performed using ImageJ.

### Statistical analysis

All statistical analyses were performed using GraphPad Prism. p values p ≤ 0.05 were considered statistically significant. (n.s. no significance, * p ≤ 0.05, ** p ≤ 0.01, *** p ≤ 0.001, **** p ≤ 0.0001). In the case of multiple comparisons, the p-values were adjusted using Bonferroni correction.

## Results

### US guided FUS systems for targeted BBB/BTB opening in GL26 glioma tumor

In this study, we employed an orthotopic GL26 glioma model in which GL26 cells expressing firefly luciferase and eGFP were stereotactically implanted in B6 Albino mice. To validate that MB-FUS enhances delivery across the BBB/BTB, we first assessed delivery of 70 kDa fluorescently labeled dextran (diameter ∼14 nm, comparable to AAV9 capsids at ∼20-25 nm) to GL26 tumors with and without FUS, administered intravenously 1 minute prior to sonication. MB-FUS was performed two weeks post-implantation using an ultrasound-guided focused ultrasound (USgFUS) system integrating 1) an imaging array (L12-5, Philips/ATL) that enabled precise spatial targeting through real-time visualization of tumor anatomy and co-registration with pre-acquired MRI and 2) a 1.5-MHz therapeutic transducer (128-element 2D array) (Fig. 1A-B, Suppl. Video 1) [44]. The therapeutic array enabled tumor-specific treatment region design through volumetric raster-scanning, with a 5 × 5 grid (0.5 mm spacing) configured to match tumor volume and achieve full coverage. Sonication was performed concurrently with intravenous MB administration (2.5 × 10^8^ MBs/kg) using the following parameters: 320 kPa peak negative focal pressure, 1 ms pulse duration for each grid point, 5 Hz pulse repetition frequency of the grid and 120 s total treatment time. During treatment, the imaging transducer was used to monitor cavitation dynamics in real-time and provided passive acoustic mapping (PAM) to localize acoustic emissions [47]. This feedback mechanism ensured sonication remained within safety thresholds.

**Fig. 1.**
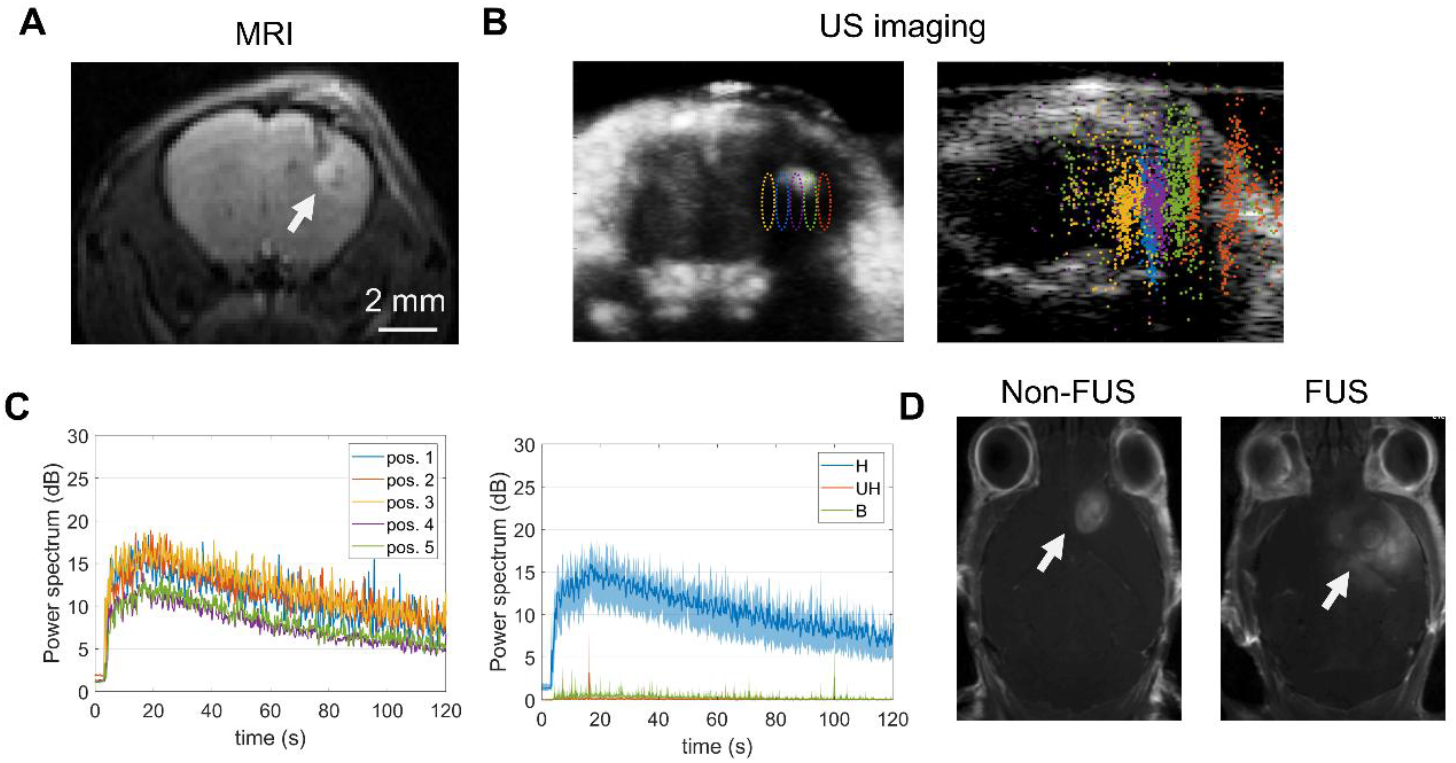
Ultrasound-guided MB-FUS treatment and BBB opening in GL26 glioma. (A) Representative T2 weighted MR image of tumor two weeks after implantation. (B) Corresponding maximum intensity projection image from contrast-enhanced ultrasound images after MB administration showing the tumor and FUS target area (left; each colored ellipse indicates a focus location from the treatment grid aligned with the imaging plane). Post-FUS passive acoustic map localizing acoustic emissions from MBs for the entire treatment (right; each colored point cloud corresponds to a focus location and each point to an individual treatment pulse). (C) Representative power levels of acoustic emissions originating from MB oscillations during the 120-s treatment. Emissions are processed in the frequency domain in specific bands (H: harmonics, UH: ultra-harmonics, B: broadband). Left: power spectrum as a function of time in the H band for each of the 5 in-plane focus locations. Right: average power spectrum for all 5 focus locations for the H, UH and B frequency bands as a function of time. (D) Representative contrast enhanced T1-weighted MR image 10 minutes after MB-FUS treatment (right) or control non-FUS treatment (left). Without FUS, contrast accumulation within the tumor results from the leaky vascular environment (left). After MB-FUS treatment (right), the wider uptake in the tumor area indicates successful BBB opening. White arrows indicate tumor locations.

Contrast-enhanced ultrasound imaging following MB administration enabled visualization of tumor morphology and location for confirmation of targeting during treatment (Fig. 1B, Suppl. Fig. 1A). Post-treatment passive acoustic mapping (PAM) analysis revealed spatially localized cavitation events during sonication, validating accurate tumor targeting (Fig. 1C). Additionally, ultra-harmonics and broadband noise remained low throughout treatment, indicating stable MB oscillation without collapse (Fig. 1C, Suppl. Fig. 1B). This is critical for safety, as broadband emissions are associated with MB destruction and potential hemorrhage. To confirm vascular barrier opening, we performed gadolinium-enhanced T1-weighted MRI 10 minutes after sonication. Focal contrast enhancement localized to the sonicated tumor region (Fig. 1D) demonstrated successful BBB/BTB opening with precise spatial targeting.

Importantly, ex vivo whole-brain fluorescence imaging confirmed 2-fold enhanced dextran accumulation (p = 0.0015) in sonicated tumors compared to untreated controls 1 hour following MB-FUS treatment (Fig. 2A-B). At the microscopic level, dextran showed substantially greater penetration and distribution throughout MB-FUS-treated tumors compared to non-FUS controls (Fig. 2C). Together, these data demonstrate that MB-FUS enhances delivery of macromolecular therapeutics to orthotopic gliomas in immunocompetent mice.

**Fig. 2.**
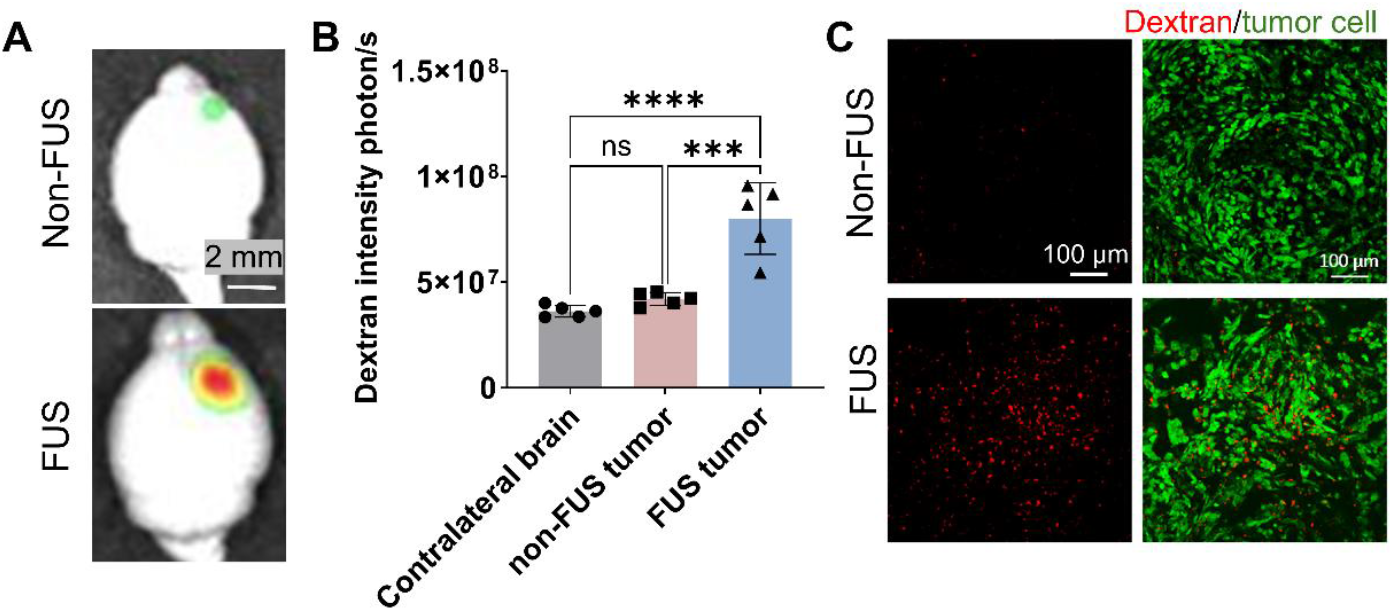
MB-FUS enhanced 70kDa Dextran delivery in GL26 glioma. (A) Representative ex vivo images of 70 kDa dextran distribution in the brain 1 hour after MB-FUS treatment (bottom) or control non-FUS treatment (top). (B) Quantification of ex vivo imaging of dextran delivery in the brain 1 hour after treatment. (C) Representative confocal microscopy of dextran distribution in the tumor. Red: dextran, green: tumor cells. p-values were determined by one-way analysis of variance (ANOVA) and were adjusted using Bonferroni correction. Plots show mean ± S.D (n=5). ***p ≤ 0.001, ****p ≤ 0.0001.

### Longitudinal PET imaging of ^64^Cu-labeled AAV9 demonstrates MB-FUS-mediated enhancement of vector delivery in GL26 glioma tumor

Having established that MB-FUS enhances macromolecular delivery to brain tumors, we next quantitatively assessed MB-FUS-mediated enhancement of AAV vector delivery in orthotopic gliomas using PET imaging of ^64^Cu-labeled AAVs. AAV9 was selected due to its well-established systemic biodistribution profile and tropism for brain tissue [29,48]. Tumor growth was monitored by MRI and bioluminescence imaging, and mice were assigned to treatment groups based on tumor volume to ensure comparable microenvironmental characteristics across groups (Fig. 3A, Suppl. Fig. 2). Two weeks post-implantation, ^64^Cu-labeled AAV9 (1×10^13^ vg/kg, 5-8 μCi/mouse) was administered intravenously immediately prior to MB-FUS treatment, following the protocol established for dextran. Real-time ultrasound imaging confirmed tumor localization (Fig. 3B), while passive acoustic monitoring demonstrated stable MB oscillations throughout treatment within the safety regime (Fig. 3C).

**Fig. 3.**
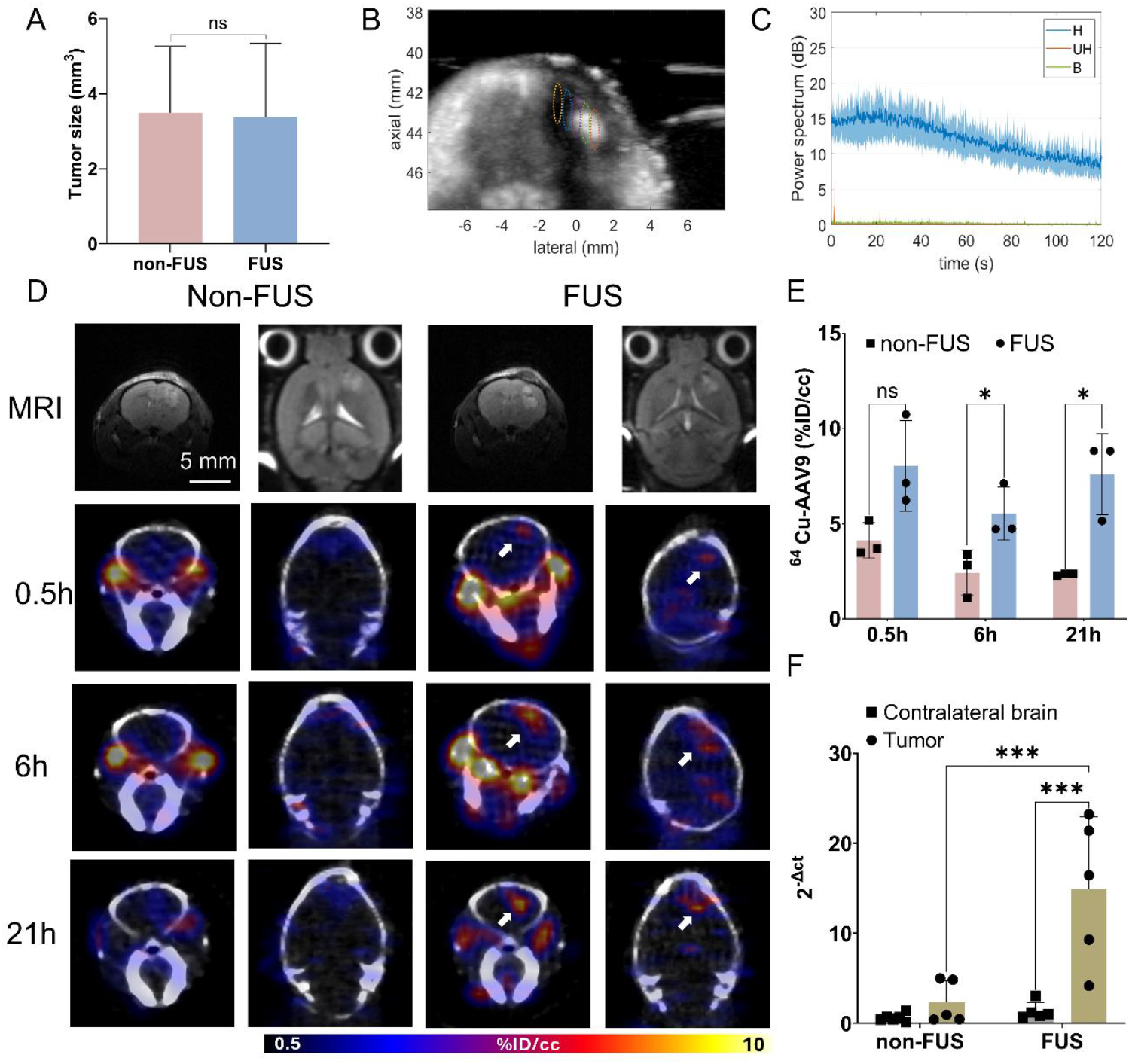
Longitudinal PET/CT imaging of ^64^Cu-labeled AAV delivery. (A) Tumor size quantification with T2 weighted MRI prior to treatment showing comparable volumes between non-FUS and MB-FUS treatment groups. (n = 5, two-tailed unpaired t test) (B) Representative MB contrast-enhanced ultrasound image with overlaid treatment grid (colored dotted lines) confirming tumor localization for targeted sonication. (C) Representative power levels of acoustic emissions originating from MB oscillations during the 120-s treatment. Emissions are processed in the frequency domain in specific bands (H: harmonics, UH: ultra-harmonics, B: broadband). (D) Representative T2-weighted MRI (top rows, acquired pre-treatment) and corresponding longitudinal PET images from the same mouse at 0.5, 6, and 21 hours post-treatment for non-FUS control (left) and MB-FUS-treated (right) mice. White arrows indicate tumor delivery. (E) Quantification of PET images in tumor regions at 0.5, 6, and 21 hours post-treatment (%ID/cc). (n = 3, two-tailed unpaired t test) (F) RT-qPCR quantification of AAV9 genome copies at 21 hours post treatment. Tumor-to-contralateral brain ratio at 21 hours demonstrating enhanced and spatially specific AAV9 delivery to FUS-treated tumors (n = 5, two-way ANOVA with Bonferroni correction). n.s. no significance, p > 0.05, *p ≤ 0.05, ***p ≤ 0.001. Data shown as mean ± SD.

To track AAV biodistribution dynamics, we performed longitudinal PET imaging at 0.5, 6, and 21 hours post-treatment. Radiolabeled capsid accumulation localized in the FUS-treated tumors, demonstrating both enhancement and spatial specificity of delivery (Fig. 3D). At the earliest timepoint (0.5 h), the ^64^Cu signal was greater in tumor regions in both groups compared to other time points, likely reflecting circulating AAVs in the vasculature. However, by 21 hours, as circulating AAVs cleared, ^64^Cu accumulation reached 7.6%ID/cc in FUS-treated tumors compared to 2.3%ID/cc in non-FUS treated tumors, representing a 3.2-fold enhancement (n=3, p=0.004, Fig. 3E). To corroborate PET imaging findings, we quantified AAV9 genome copies in tumor tissue at 21 hours using qPCR, which showed a 6.4-fold enhancement in FUS-treated tumors compared to non-treated tumors (n=5, p=0.0003, Fig. 3F), confirming the PET imaging quantification.

Together, these data demonstrate that MB-FUS significantly enhances spatially targeted AAV9 delivery to orthotopic gliomas with sustained vector retention. Moreover, quantitative ^64^Cu-PET imaging provides non-invasive assessment of AAV biodistribution and pharmacokinetic insights that can inform optimization of treatment timing and treatment parameters for future therapeutic applications.

### Enhanced AAV delivery with MB-FUS resulted in higher transduction efficiency in GL26 glioma tumor

To assess whether enhanced AAV delivery can lead to functional transgene expression, we administered AAV9 encoding the tdTomato fluorescent reporter and evaluated transduction efficiency in tumors. Given the aggressive growth of the GL26 tumors and the timeline for AAV transduction (peak expression at approximately 1-month post-administration), we modified the treatment protocol to perform treatment 3 days post tumor implantation, providing a sufficient window to monitor transgene expression before animals reached endpoint.

Strong harmonic emissions with minimal broadband emissions occurred throughout the 120-second sonication period (Fig. 4A), consistent with safe cavitation dynamics observed in previous experiments. Notably, harmonic emission levels were correlated with subsequent tdTomato expression intensity (Fig. 4A-B, Suppl. Fig.3), suggesting that acoustic monitoring can serve as a real-time predictor of treatment effectiveness and transduction outcomes.

**Fig. 4.**
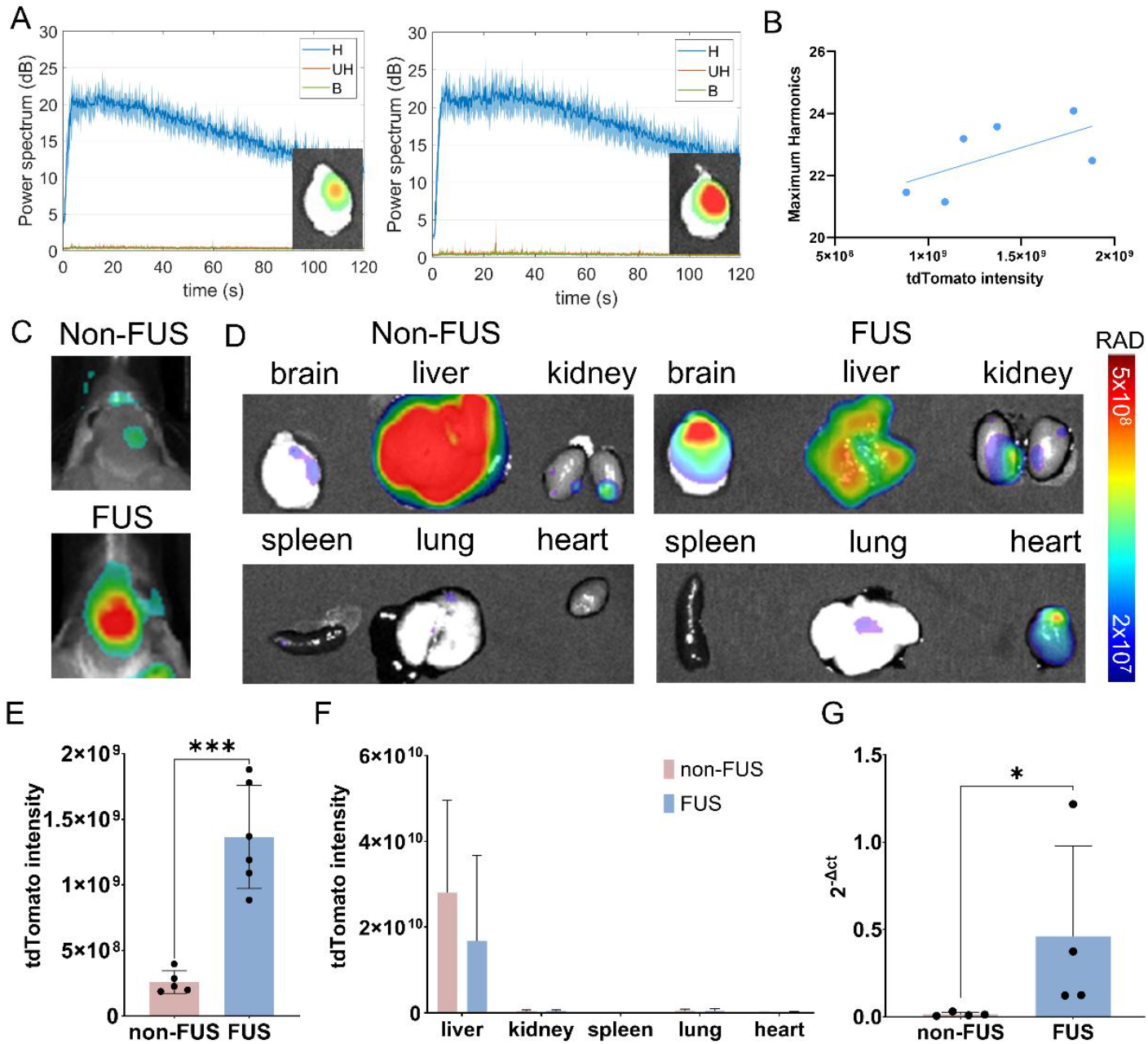
AAV transgene expression in brain tumors and peripheral organs following MB-FUS treatment. (A) Representative acoustic emission spectra during treatment (H: harmonics; UH: ultraharmonics; B: broadband) with endpoint bioluminescence images (insets). (B) Correlation between harmonic levels and tdTomato expression. (C) Representative in vivo tdTomato fluorescence at 14 days post-treatment. (D) Ex vivo organ biodistribution showing AAV9 tropism for non-FUS AAV9 only (left) and FUS group (right). (E) Quantification of ex vivo tdTomato intensity in the brain tumor region (n = 5-6). (F) Peripheral organ fluorescence quantification. (n = 5-6) (G) qPCR validation of tumor transgene expression (n=4). p-values were determined by two-tailed unpaired t test. *p ⩽ 0.05, ***p ⩽ 0.001. Data shown as mean ± SD.

Robust tdTomato expression was localized to the MB-FUS-treated tumor region at 17 days post-treatment (Fig. 4C), with a 5.3-fold increase in tdTomato fluorescence intensity in MB-FUS-treated tumors compared to non-FUS-treated tumors (p=0.0002, Fig. 4E). Thus, enhanced AAV delivery translated to significantly increased transgene expression. Biodistribution analysis of major organs showed expected AAV9 tropism, with high expression in the liver for both groups (Fig. 4D). Interestingly, a trend toward reduced hepatic tdTomato expression was observed in MB-FUS-treated mice, although not statistically significant (Fig. 4F), potentially reflecting more efficient brain retention of systemically administered AAV. Quantification of tdTomato mRNA levels by RT-qPCR in tumor tissue corroborated the optical imaging findings, demonstrating significantly elevated transgene expression in MB-FUS-treated tumors (Fig. 4G, p = 0.029).

To examine the spatial distribution and cellular localization of AAV transduction within the tumor microenvironment, we performed fluorescence microscopy on tumor tissue sections at 17 days post-treatment. In non-FUS-treated tumors, minimal tdTomato-positive cells were observed, with only sparse transduction events within the tumor (Fig. 5, top row). In contrast, tdTomato expression was greater in MB-FUS-treated tumors, with widespread transduction throughout the tumor tissue (Fig. 5, bottom row). Notably, tdTomato signal colocalized with eGFP-positive tumor cells in the merged images, confirming successful transduction of tumor cells.

**Fig. 5.**
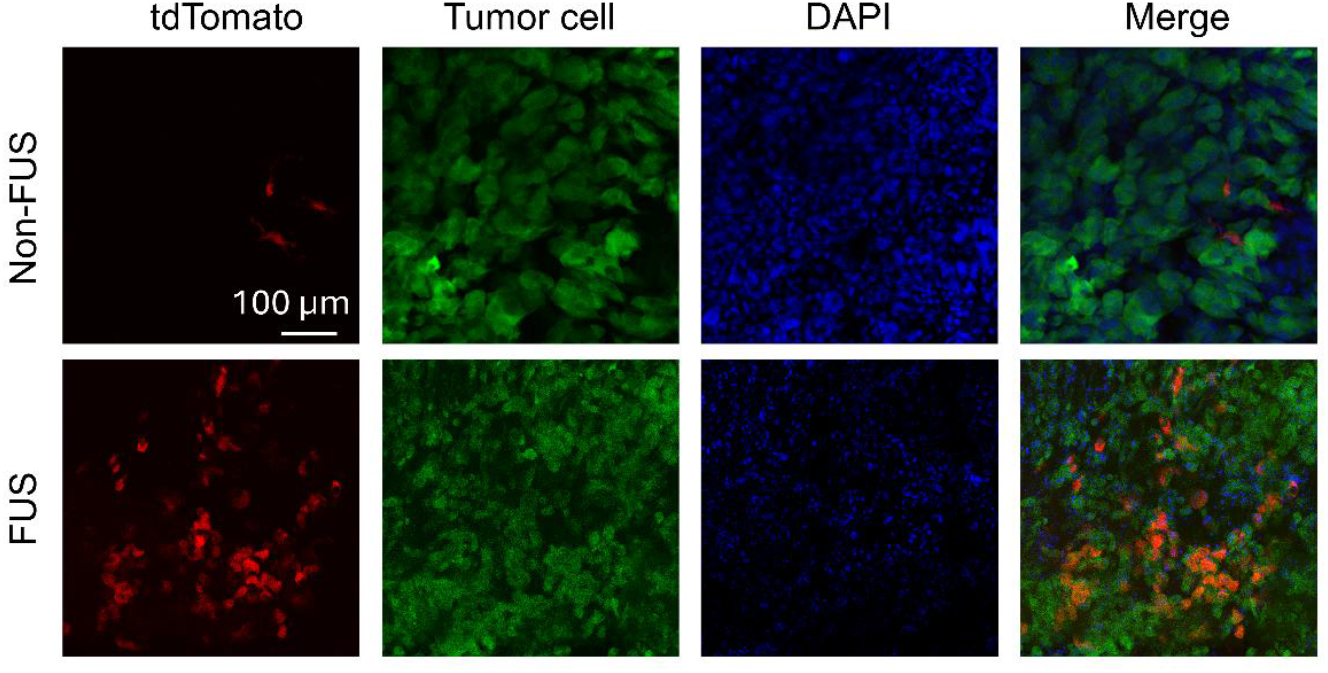
Microscopy of enhanced AAV transgene expression in tumor tissue following MB-FUS. Representative fluorescence microscopy of tumor sections at 14 days post-treatment. Red: tdTomato (AAV transgene); Green: eGFP (tumor cells); Blue: DAPI (nuclei); Merge: overlay showing colocalization. Non-FUS control (top) shows minimal transduction, while FUS-treated tumor (bottom) demonstrates extensive tdTomato expression throughout tumor tissue with substantial tumor cell colocalization. Images are acquired and presented as same scale Scale bar: 100 µm.

Collectively, MB-FUS treatment achieved enhanced functional transgene expression in gliomas with improved tumor cell transduction, demonstrating that increased AAV delivery translates to functional efficacy, while acoustic monitoring provided real-time prediction of treatment outcomes.

## Discussion

In this study, we demonstrated that microbubble-enhanced focused ultrasound (MB-FUS) significantly enhances systemic AAV delivery to orthotopic GL26 gliomas and that quantitative PET imaging of radiolabeled AAV provides a method to gauge the vector biodistribution, yielding critical pharmacokinetic insights for optimizing treatment protocols. Using an ultrasound-guided FUS platform integrated with real-time acoustic monitoring, we achieved spatially precise AAV delivery that translated to robust transgene expression in the sonicated tumor regions. These findings establish MB-FUS as a promising non-invasive strategy to overcome BBB/BTB limitations for AAV-mediated gene therapy in brain cancers.

We found that MB-FUS can enhance AAV delivery to brain tumors and quantitative PET imaging provides absolute measurements of AAV biodistribution following MB-FUS treatment. Previous studies have primarily relied on indirect measures of BBB disruption, such as gadolinium-enhanced MRI, which confirm barrier opening but do not quantify therapeutic agent delivery. While gadolinium extravasation demonstrates successful BBB modulation, the pharmacokinetics of large biologics such as AAV vectors differ substantially from small contrast agents [42]. Our ^64^Cu-PET imaging approach directly measures AAV biodistribution in absolute units (%ID/cc) [44,46], achieving 7.6%ID/cc tumor delivery with FUS treatment (3.2-fold increase over controls at 21 hours) while providing temporal pharmacokinetic data on delivery kinetics and retention. While both treated and control tumor regions showed elevated signal at early timepoints due to circulating AAVs, the differential signal between MB-FUS-treated and non-treated tumors was increasingly pronounced over time, indicating both enhanced initial extravasation and sustained retention in sonicated regions. This quantitative pharmacokinetic information is critical for rational optimization of treatment parameters, including optimal timing between AAV administration and sonication, sonication duration, and combination strategies with other therapies.

The ultrasound-guided FUS platform employed in this study enabled precise spatial targeting through real-time visualization of tumor anatomy. The phased-array transducer provides electronic beam steering, allowing tumor-specific treatment grid design, a capability critical for clinical translation, where patient-specific tumor morphology and location require customized treatment planning. Real-time feedback on microbubble cavitation dynamics resulted from passive acoustic mapping (PAM) throughout treatment, ensuring robust treatment while remaining within safety limits. Notably, we observed a positive correlation between harmonic emission levels during treatment and subsequent transgene expression, suggesting that acoustic parameters may serve as predictive metrics for treatment efficacy and enable adaptive protocols that optimize outcomes in real-time. Integration of acoustic monitoring with previously demonstrated closed-loop control systems [49–51], could be extended to optimize AAV gene therapy delivery through further investigation of acoustic-transduction relationships.

Critically, MB-FUS-enhanced AAV delivery translated to robust functional transgene expression, with 5.3-fold increased transduction in treated tumors. Widespread tdTomato expression throughout the tumor parenchyma colocalized in tumor cells, confirming efficient AAV penetration beyond the perivascular space into the tumor microenvironment. The widespread transduction pattern suggests MB-FUS enhances AAV penetration beyond perivascular regions into deep tumor tissue, potentially overcoming elevated interstitial pressure and the restrictive tumor microenvironment. While this study employed tdTomato as a reporter, the platform is readily translatable to therapeutic transgenes targeting glioma treatment resistance, including immunomodulatory genes (IL-12, IFN-β), tumor suppressors (p53, PTEN), pro-apoptotic factors (TRAIL), and anti-angiogenic agents. Furthermore, recent advances in capsid engineering have generated AAV variants with enhanced tumor tropism [52–55]. Combining these tumor-targeted capsids with MB-FUS could yield synergistic improvements in both delivery efficiency and cellular specificity.

## Conclusion

This study establishes MB-FUS as an effective strategy to enhance AAV delivery to brain tumors and demonstrates that quantitative PET imaging enables treatment monitoring and optimization. This integrated platform, combining targeted delivery, predictive acoustic monitoring, and pharmacokinetic quantification, provides essential tools for developing optimized gene therapies for brain malignancies.

## Data availability

All data supporting the findings of this study are available within the article and its supplementary files. Any additional requests for information can be directed to, and will be fulfilled by, the corresponding authors.

## Acknowledgments

We thank Dr. Frezghi Habte, Dr. Edwin Chang, and Laura Jean Pisani at the Stanford Center for Innovation in In Vivo Imaging (SCi3) for providing facilities for orthotopic tumor inoculation and technical support with PET/CT, optical imaging, and MRI studies. This study was supported by NIH grants R01 CA112356, R01 EB028646 and the Focused Ultrasound Foundation. Y.G. was partially supported by the Focused Ultrasound Foundation Lockhart Postdoctoral Fellowship.

## Author contributions

Y.G. and K.W.F. designed research; Y.G, J.F., and N.Z. performed FUS treatment; J.W.S. performed AAV radio labeling; Y.G, N.Z., J.W.S., J.W., and B.L.J. performed PET/CT imaging; Y.G, B.L.J. and M.N.R. inoculated tumors; Y.G.; S.K.T cultured tumor cells; Y.G. and S.K.T performed the bioassay and analyzed the data, B.L.J. and M.N.R. performed animal care, Y.G. and K.W.F. wrote the manuscript. All authors reviewed the manuscript and approved the final version.

## Competing interests

The authors declare that they have no competing interests.

## Notes

### Competing Interest Statement

The authors have declared no competing interest.

### Summary of Updates

updated method section on Radiolabeling of AAV; Supplementary Video added

