## Supplementary Information for "Microbubble-Enhanced Focused Ultrasound Improves Targeted Adeno-Associated Virus Delivery in Brain Tumors Quantified by PET Imaging"

**Affiliations**

**Supplementary Information**

**Supplementary Figures**

**Suppl. Fig. 1.** A) Maximum intensity projection from contrast-enhanced ultrasound imaging demonstrating microbubble flow through a tube phantom with overlaid focal zone markers (colored dots), confirming precise FUS targeting capability. (B) Representative passive acoustic spectrogram acquired during MB-FUS sonication of a GL26 tumor-bearing mouse.

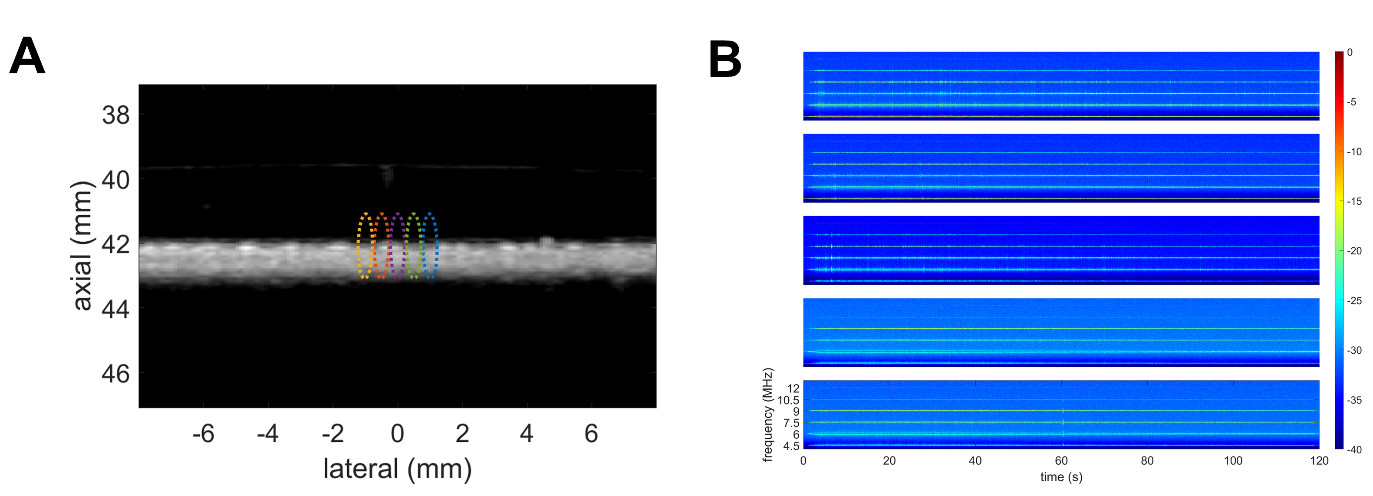

**Suppl. Fig. 2.** A) Microplate reader (Spark, Tecan Life Science) quantification confirming eGFP fluorescence and firefly luciferase bioluminescence stably transduced in GL26 cells (mean ± SD). B) Timeline for tumor implantation and monitoring (upper) and longitudinal bioluminescence imaging (lower) showing tumor growth over two weeks post-inoculation with representative optical/MRI images and fluc intensity quantification.

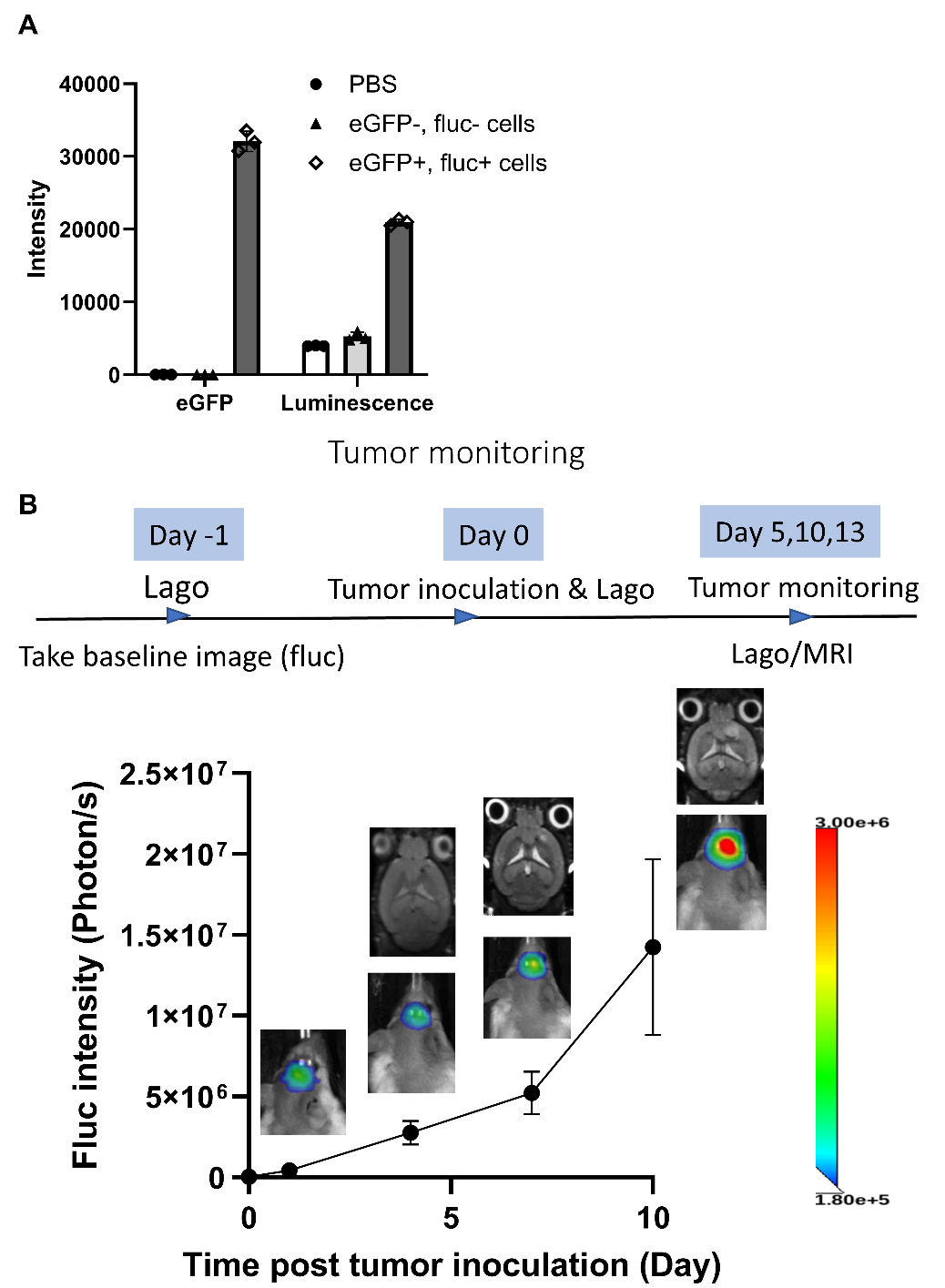

**Suppl. Fig. 3.** Correlation between tdTomato fluorescence intensity at 17 days post-treatment and (A) harmonic, (B) ultraharmonic, and (C) broadband emission levels measured during MB-FUS sonication.

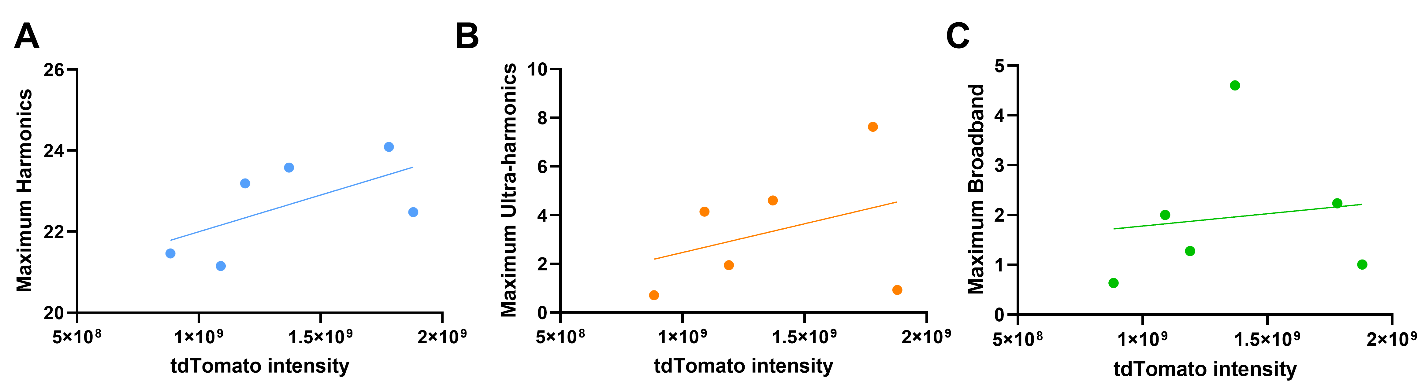
